# Auranofin inhibits virulence in *Pseudomonas aeruginosa*

**DOI:** 10.1101/198820

**Authors:** Leon Zhen Wei Tan, Joey Kuok Hoong Yam, Ziyan Hong, May Margarette Santillan Salido, Bau Yi Woo, Sam Fong Yau Li, Liang Yang, Michael Givskov, Shu-Sin Chng

## Abstract

*Pseudomonas aeruginosa* is widely attributed as the leading cause of hospital-acquired infections. Due to intrinsic antibiotic resistance mechanisms and the ability to form biofilms, *P. aeruginosa* infections are challenging to treat. *P. aeruginosa* employs multiple virulence mechanisms to establish infections, many of which are controlled by the global virulence regulator Vfr. An attractive strategy to combat *P. aeruginosa* infections is thus the use of anti-virulence compounds. Here, we report the discovery that FDA-approved drug auranofin attenuates virulence in *P. aeruginosa*. We demonstrate that auranofin acts by targeting Vfr, which in turn leads to inhibition of quorum sensing (QS) and Type IV pili (TFP). Consistent with inhibition of QS and TFP expression, we show that auranofin attenuates biofilm maturation, and when used in combination with colistin, displays strong synergy in eradicating *P. aeruginosa* biofilms. Auranofin may have immediate applications as an anti-virulence drug against *P. aeruginosa* infections.

## Introduction

*Pseudomonas aeruginosa* is a prevalent opportunistic pathogen known for its potent virulence, environmental versatility and multi-drug resistance. Its role as the leading cause of hospital-acquired infections (HAIs) among immuno-compromised patients is well documented^1^. Notably, it is one of the main causes of pneumonia in cystic fibrosis (CF) patients^2^, and wound infections in surgical patients or burn victims^3^. Furthermore, *P. aeruginosa* has been known to thrive in a variety of ecological niches^4^, allowing colonization on abiotic surfaces like catheters and other implanted medical devices^5,6^. Increasing incidences of multi-drug resistant (MDR) *P. aeruginosa* infections^7,8^ in the clinic pose a serious threat to global health, underscoring the need to develop new strategies to combat these pathogens.

*P. aeruginosa* carries several intrinsic antibiotic resistance mechanisms involving the collaboration of restricted uptake through the outer membrane^9^, the use of energy-dependent efflux pumps^10,11^, and expression of β-lactamases^12,13^. In addition, *P. aeruginosa*, like many other bacteria, exhibits a form of social behavior known as biofilm formation^14^, which further increases its tolerance to antibiotics^15^. A biofilm is a community of bacteria encased in a polymeric mesh consisting of extracellular DNA (eDNA), proteins and polysaccharides. In the clinic, significantly higher patient mortality has been associated with the formation of biofilms in the lungs of CF patients^16^. Specifically, the presence of biofilms increases the complexity of treatment due to the function of the extracellular matrix as a physical barrier against, and sponge for, antimicrobial drugs^15^. The small pore size of the biofilm mesh effectively reduces penetration of lysozymes and macromolecular drugs, presumably decreasing the immediate concentrations of these agents surrounding the cells. Additionally, the biofilm matrix is also highly negatively charged due to the presence of eDNA and polysaccharides, which protect the colony from positively-charged aminoglycosides^17^. Consequently, the inhibition of biofilm formation, maturation and/or function remains a principal consideration in the search for new treatment options against *P. aeruginosa*.

An attractive alternative strategy to control bacterial infections has been the use of small molecules that block virulence. Such anti-virulence compounds may reduce the ability of bacteria to infect the host^18,19^, and/or attenuate bacterial defenses against host immune cells^20^, thereby facilitating clearance of the infection site^21^. The virulence of *P. aeruginosa* is multi-faceted but largely controlled by the global virulence factor regulator, Vfr^22^. Vfr is a cyclic adenosine monophosphate (cAMP)-dependent transcription factor that coordinates expression of roughly 200 genes^23,24^. It regulates virulence directly by activating expression of secretion systems^25^ (e.g. Type III) and the pilus formation machinery^26^ (Type IV). The Type III secretion system (T3SS) in *P. aeruginosa* mediates cytotoxicity against host cells through the delivery of molecular effectors that can disrupt host processes^27^. The Type IV Pilus (TFP) is important for twitching motility^28^, and often associated with the establishment and maturation of *P. aeruginosa* biofilms. Vfr also controls quorum sensing (QS)^29^, a cell-cell communication network involving the *las-, rhl-* and *pqs*-encoded systems, which plays a role in the production of several virulence factors including extracellular proteases, biosurfactants and toxins^20^, and is important for biofilm maturation^30^ and immune evasion. While many small molecules that inhibit QS and T3SS have been reported^31-33^, no inhibitors of Vfr and TFP have been discovered to date.

In this work, we identified auranofin, an FDA-approved, gold(I)-based anti-inflammatory drug, as a potent inhibitor of virulence in *P. aeruginosa*. We show that auranofin inhibits QS and TFP formation at low micromolar concentrations. It does this in part by targeting cysteines in Vfr and abolishing the ability of Vfr to bind DNA. Using flow cell biofilm assays, we demonstrate that auranofin is able to attenuate biofilm maturation. Furthermore, auranofin displays strong synergistic effects in eradicating *P. aeruginosa* biofilms *in vitro*, when used in combination with colistin. This synergy is also observed in a murine infection model. We propose that auranofin can potentially be re-purposed as an anti-virulent drug to treat *P. aeruginosa* infections.

## Results

### Auranofin inhibits QS in *Pseudomonas aeruginosa*

We initially discovered auranofin in a screen to identify small molecules that inhibit QS in *P. aeruginosa*. We screened ∼6,000 compounds from various libraries using a *lasB-gfp*(ASV) strain^34^, where expression of a fusion construct between LasB N-terminus and an unstable GFP variant (GFP(ASV)) is controlled by constitutively expressed LasR, the cognate transcriptional regulator of the *las* pathway. Inhibitors of this pathway should reduce GFP fluorescence in this assay. In addition, since the *las* pathway controls both the *rhl* and *pqs* pathways, compounds identified in this screen could potentially shut down the entire QS network. We found that auranofin exhibits sub-micromolar (IC_50_) inhibition of LasB-GFP expression with minimal effects on cell growth (Fig. 1A and S1). Auranofin also strongly inhibits fluorescence in two other reporter strains, *rhlA-gfp*(ASV) and *pqsA-gfp*(ASV)^35^, both under the control of their respective transcriptional activators, RhlR and PqsR (Fig. 1B and 1C). To validate our findings, we examined the effects of auranofin on the levels of RNA transcripts for key QS-regulated genes, as well as on the production of various QS-controlled virulence factors. Auranofin results in the down-regulation of *lasA* and *rhlA* transcripts (Fig. S2), and significantly reduces the levels of elastase (LasB), rhamnolipids and pyocyanin (Fig. 2). Taken together, these results establish that auranofin is a potent QS inhibitor.

**Figure 1.**
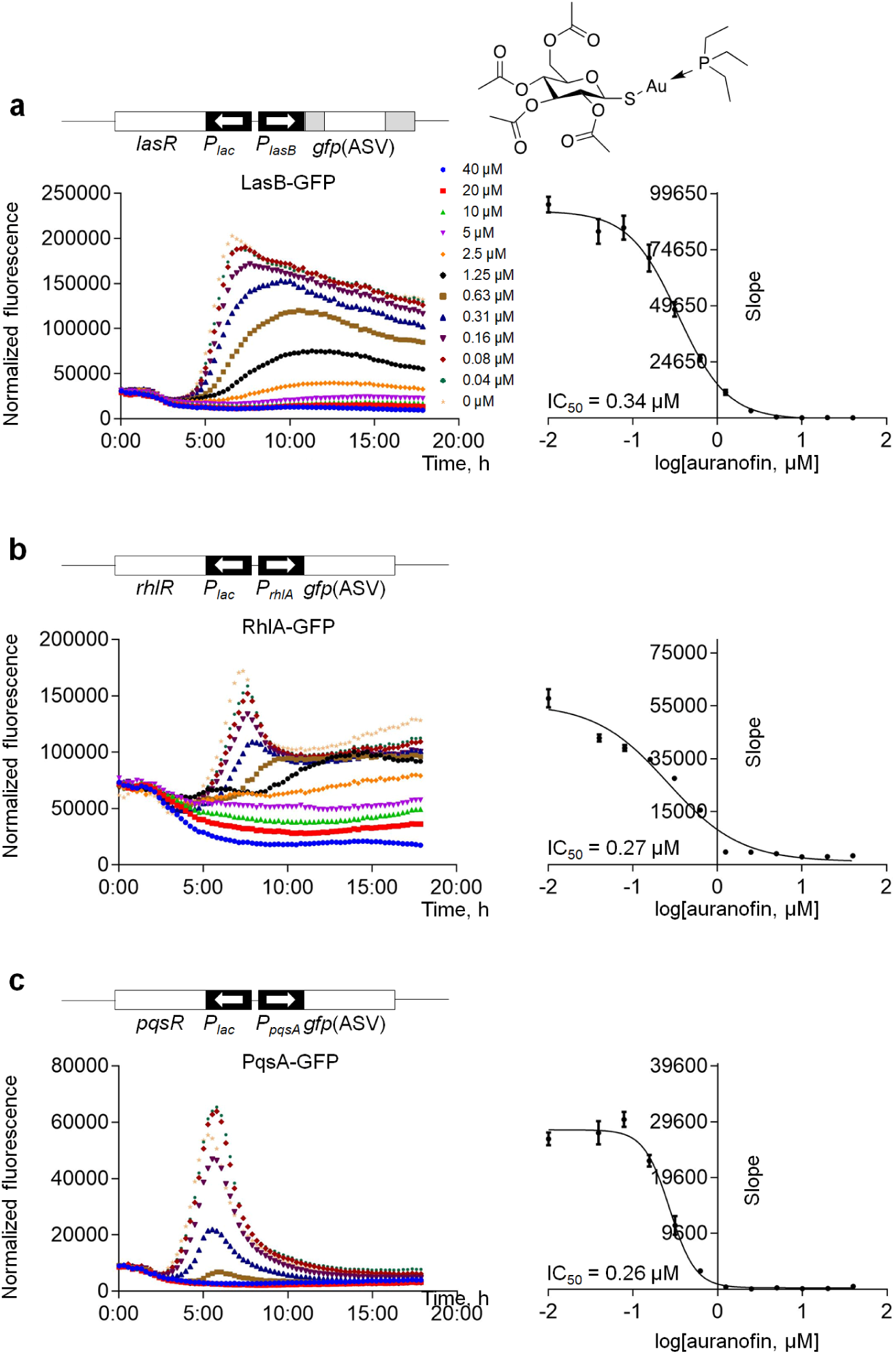
Auranofin inhibits all levels of QS with minimal effects on cell growth. Representative fluorescence induction profiles (relative fluorescence units normalized to cell density, over time) for (a) LasR-mediated expression of LasB-GFP, (b) RhlR-mediated expression of P_*rhlA*_-*gfp*, and (c) PqsR-mediated expression of P_*pqsA*_-*gfp*, in the presence of indicated concentrations of auranofin, whose chemical structure is shown on the top. Dose-dependent inhibition curves for each assay are plotted on the right. Error bars represent standard deviation from technical triplicates.

**Figure 2.**
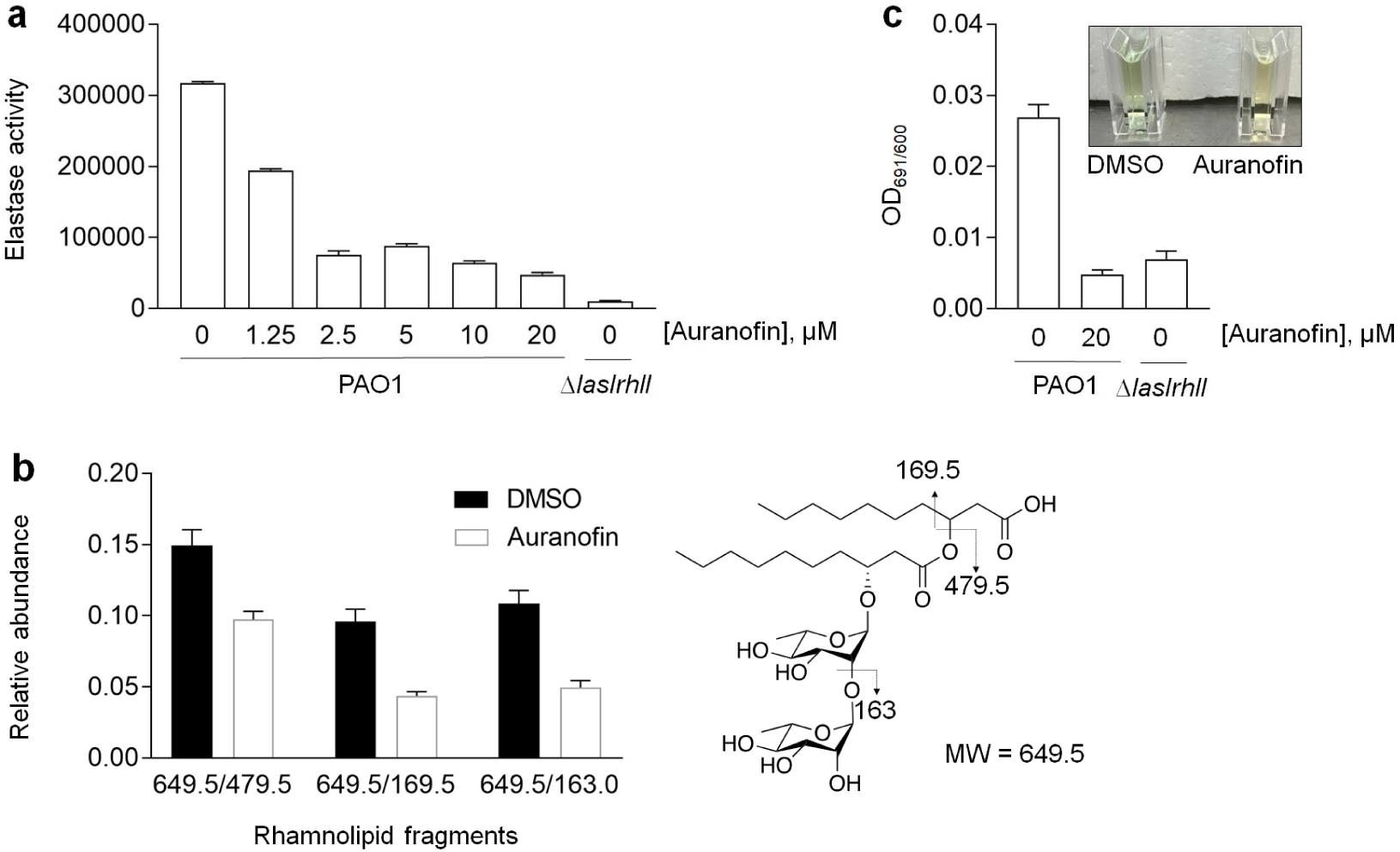
The production of key QS-controlled virulence factors is reduced in the presence of auranofin. (a) Elastase activity in the supernatants of PAO1 and Δ*lasIrhlI* cultures treated with indicated concentrations of auranofin. (b) Relative abundance of rhamnolipid fragments isolated from PAO1 cells treated with 20 µM auranofin, as quantified using multiple reaction monitoring mass spectrometry. The chemical structure of rhamnolipid and its measured fragments are shown on the right. (c) Relative levels of pyocyanin in supernatants of PAO1 and Δ*lasIrhlI* cultures treated with indicated concentrations of auranofin, as measured at absorbance at 691 nm. Error bars represent standard deviation from biological triplicates.

### Auranofin inhibits expression of Type IV pili (TFP), and likely Type III secretion system (T3SS), in *P. aeruginosa*

To gain insights into the mechanism(s)-of-action of auranofin, we compared the protein profiles of *P. aeruginosa* in the presence and absence of auranofin. Proteomic analysis showed that about 210 proteins were significantly up- or down-regulated in the presence of auranofin. Several efflux pumps were the most up-regulated, while certain outer membrane porins were down-regulated in the presence of auranofin (Table S1). These changes indicate a response to prevent intracellular build-up of auranofin. Notably, consistent with it being a QS inhibitor, treatment with auranofin caused reduction in the levels of proteins involved in or regulated by QS. Interestingly, we also observed down-regulation of proteins involved in T3SS and TFP formation. We showed that auranofin is able to inhibit TFP-dependent motility, including swarming and twitching^36,37^ (Fig. 3). Our observations here are, in part, due to transcriptional down-regulation of these systems (Fig. S2). These results imply that auranofin either has multiple targets, and/or it targets a transcriptional regulator upstream of QS, T3SS and TFP.

**Figure 3.**
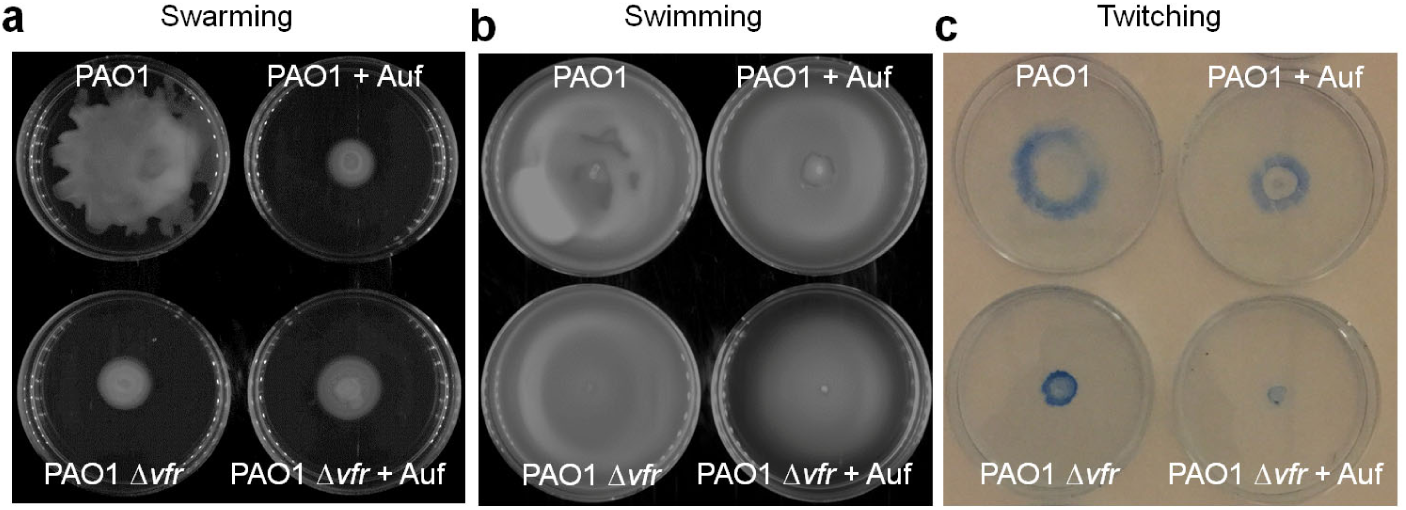
Motility phenotypes that depend on TFP (i.e. swarming and twitching) are reduced in the presence of auranofin. (a) Swarming (0.5% LB agar), (b) swimming (0.3% LB agar), and (c) twitching (1% LB agar) motilities of PAO1 and PAO1 Δ*vfr* strains in the presence or absence of 20 µM auranofin. For twitching, cells at the base of the plates were stained with Coomassie blue.

### Auranofin inhibits Vfr *in vitro* but has other targets in cells

QS, T3SS and TFP formation are positively regulated by Vfr^24^. We asked if auranofin directly targets Vfr function. To test this, we examined the ability of auranofin to inhibit the function of Vfr in binding to promoter sequences *in vitro* using electrophoretic mobility shift assays (EMSA). We first showed that purified Vfr is able to specifically bind DNA fragments containing the promoter region of *lasR*, a representative QS gene (Fig. S3), as previously reported^38^. We further demonstrated that auranofin effectively prevents Vfr binding to this promoter in a dose-dependent manner, indicating that it inhibits Vfr function *in vitro* (Fig. 4A). Gold(I)-based compounds are thiophilic, and auranofin has previously been shown to react covalently with cysteine residues within target proteins^39,40^. To ascertain if its inhibition of Vfr proceeded via covalent modification of cysteines, we generated a cysteine-less Vfr variant (VfrΔ5) in which all five cysteines were replaced by serine residues. This variant is functional as judged by its ability to bind the *lasR* promoter *in vitro* (Fig. S3). Remarkably, auranofin is no longer able to prevent binding of VfrΔ5 to this promoter (Fig. 4B). These results establish that auranofin inhibits Vfr *in vitro* via its cysteine residues.

**Figure 4.**
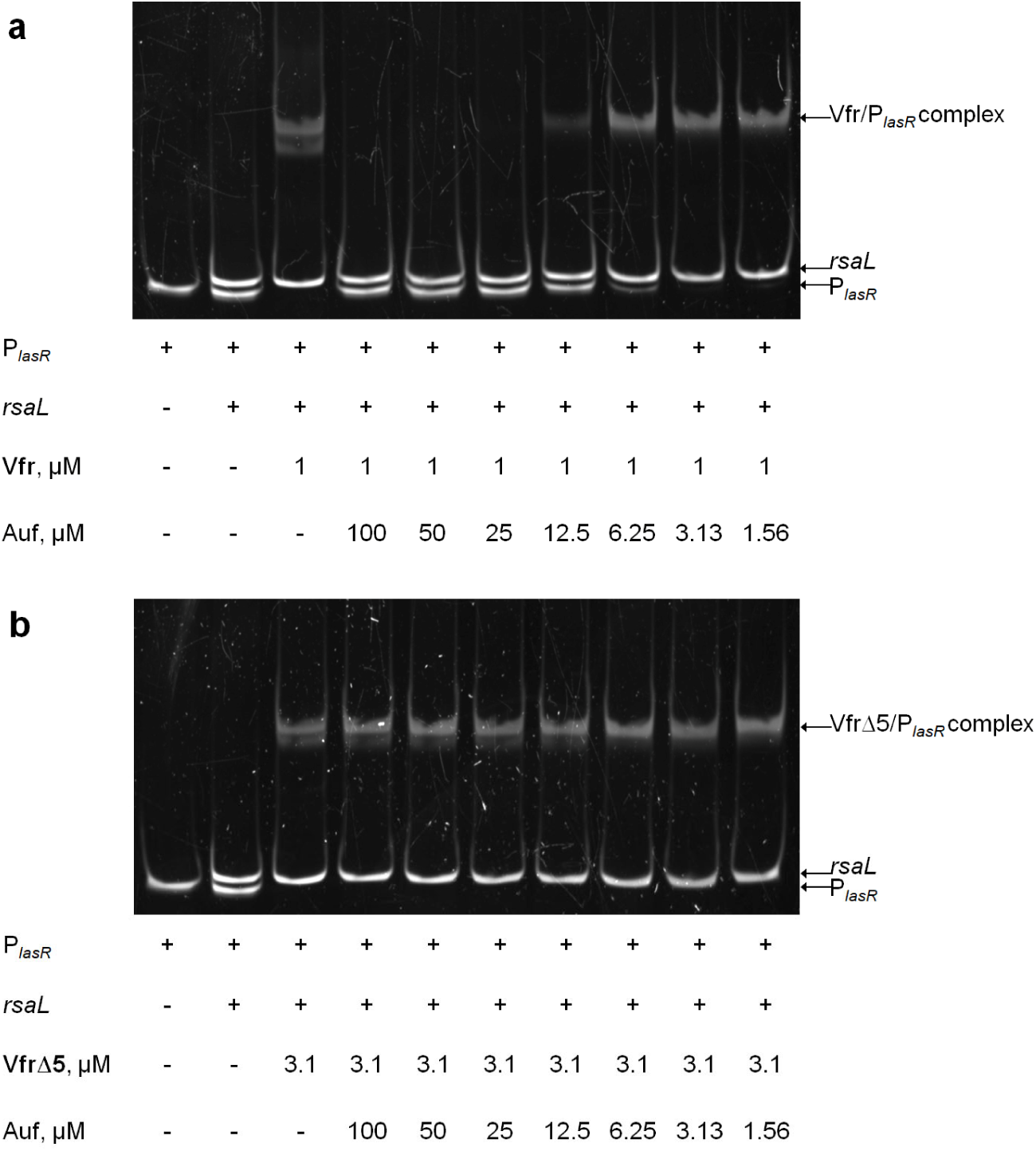
Auranofin inhibits Vfr binding to target promoter sequences *in vitro* by reacting with cysteine residues. Representative polyacrylamide gels showing specific binding of (a) Vfr and (b) VfrΔ5 to P_*lasR*_, in the presence of indicated concentrations of auranofin. Vfr and VfrΔ5 bind specifically to the target promoter sequence at ∼1 and 3 µM, respectively (see Fig. S3). The coding region of the gene *rsaL* is used as a probe to control for non-specific binding.

To determine if Vfr is the main target of auranofin in cells, we complemented the Δ*vfr* strain with wild-type Vfr and the VfrΔ5 variant, and examined the elastase levels and twitching motility in the presence and absence of auranofin. Both wild-type Vfr and the VfrΔ5 variants were able to complement the Δ*vfr* strain in these assays (Fig. S4). We showed that auranofin inhibits both phenotypes in the wild-type Vfr complemented strain, as expected. Surprisingly, auranofin was still able to inhibit these phenotypes in the VfrΔ5 complemented strain, unlike what we have observed *in vitro*. While we have demonstrated that auranofin does inhibit Vfr via its cysteine residues *in vitro*, our data suggests auranofin may have other targets, in addition to Vfr.

### Auranofin impedes the development of *P. aeruginosa* biofilms

Both TFP and QS are required during biofilm development^41,42^. TFP are important for initial surface attachment of cells, as well as for the organization of cells into the typical “mushroom” architecture of mature *P. aeruginosa* biofilms. QS is also critical for biofilm maturation, although the mechanisms here are less understood. Since auranofin inhibits both TFP formation and QS networks, we expect it to also have an impact on the development of *P. aeruginosa* biofilms. To test this, we monitored biofilm formation and maturation in the presence and absence of auranofin over a period of 72 h in continuous flow chambers, using *ΔlasIΔrhlI* as the control strain. The formation of fully matured, canonical mushroom-shaped biofilm structures for wild-type *P. aeruginosa* was readily observed within 48 h (Fig. 5A). Interestingly, while auranofin did not have obvious effects on the initial surface attachment of cells at early time points (6 h and 24 h), it impeded the development of mushroom-shaped structures beyond 48 h, and only allowed formation of microcolonies. Biofilms formed in the presence of auranofin appeared flatter, similar to biofilms of QS-null (e.g. *ΔlasIΔrhlI*) and/or TFP-null strains^43^. Detailed analyses of these biofilms revealed that biomass (Fig. 5B), mean biomass thickness (Fig. 5C), and the roughness coefficient (Fig. 5D) were all significantly reduced in the presence of auranofin after 48 h. Our data clearly indicate that despite permitting biofilm attachment and growth, auranofin attenuates the progression of biofilm maturation, likely by inhibiting QS and TFP.

**Figure 5.**
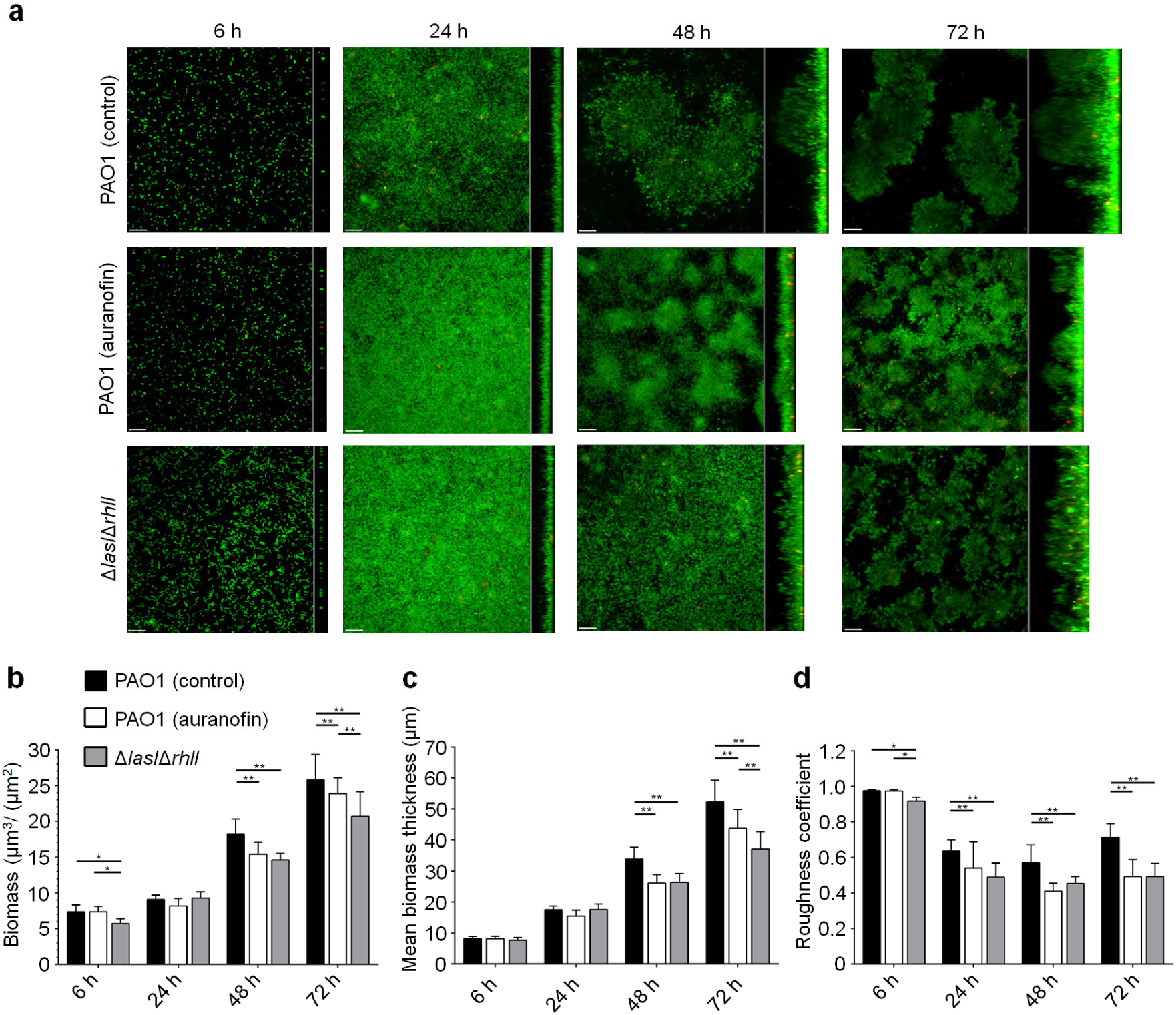
Auranofin attenuates the formation of mushroom-shaped structures in *Pseudomonas aeruginosa* biofilms. (a) Confocal fluorescence microscopy images illustrating the development of wild-type *P. aeruginosa* PAO1 biofilms in the presence or absence of auranofin over 72 h in flow chambers. QS-defective mutant strain PAO1 Δ*lasI*Δ*rhlI* was used as control. Biofilms were stained with SYTO9 before imaging. Biological replicates were performed, and a representative image for each condition/time point is shown. Scale bar, 20 µm. Biofilm parameters including (b) biomass, (c) mean biomass thickness, and (d) roughness coefficient, were quantified via analyses of a series of confocal images using COMSTAT2 analysis software. Error bars represent standard deviations from biological triplicates. Student’s t-test: *P < 0.05 and **P < 0.01.

### Auranofin exhibits synergistic effects with colistin in killing *P. aeruginosa* biofilms

Since auranofin affects the maturation of the biofilm, we asked whether it may display synergy with colistin, a last resort drug commonly used to treat *P. aeruginosa* infections. We first evaluated the combined effects of auranofin and colistin by monitoring viability of *P. aeruginosa* cells, following treatment of biofilms (pre-formed for one day on glass beads) with different concentrations of the two drugs in a checkerboard format. The highest concentrations of auranofin (10 µM) or colistin (10 µg ml^-1^) single treatment only reduced the average number of viable bacterial cells per glass bead from ∼1 x 10^7^ c.f.u. (no treatment) to ∼3.1 x 10^6^ c.f.u. or ∼5.7 x 10^6^ c.f.u., respectively (Fig. 6A). Remarkably, the number of viable bacterial cells was drastically reduced (average ∼178 c.f.u./glass bead) when the biofilms were treated with a combination of lower concentrations of auranofin (2 µM) and colistin (2 µg ml^-1^), demonstrating strong synergistic effects. We also tested the synergy of auranofin and colistin using *P. aeruginosa* flow cell biofilms. As previously reported, colistin treatment alone (at 10 µg ml^-1^) resulted in a sharp decline in cell viability across a 2-day old biofilm, with the exception of several surviving colistin-tolerant subpopulations^43^ (Fig. 6B). Interestingly, adding auranofin (1 µM) prior to biofilm formation allowed subsequent colistin treatment to completely eradicate the *P. aeruginosa* biofilm. In addition, co-treatment of a 2-day biofilm with auranofin and colistin also resulted in complete killing. Taken together, these results demonstrate strong synergistic effects between auranofin and colistin against *P. aeruginosa* biofilms.

**Figure 6.**
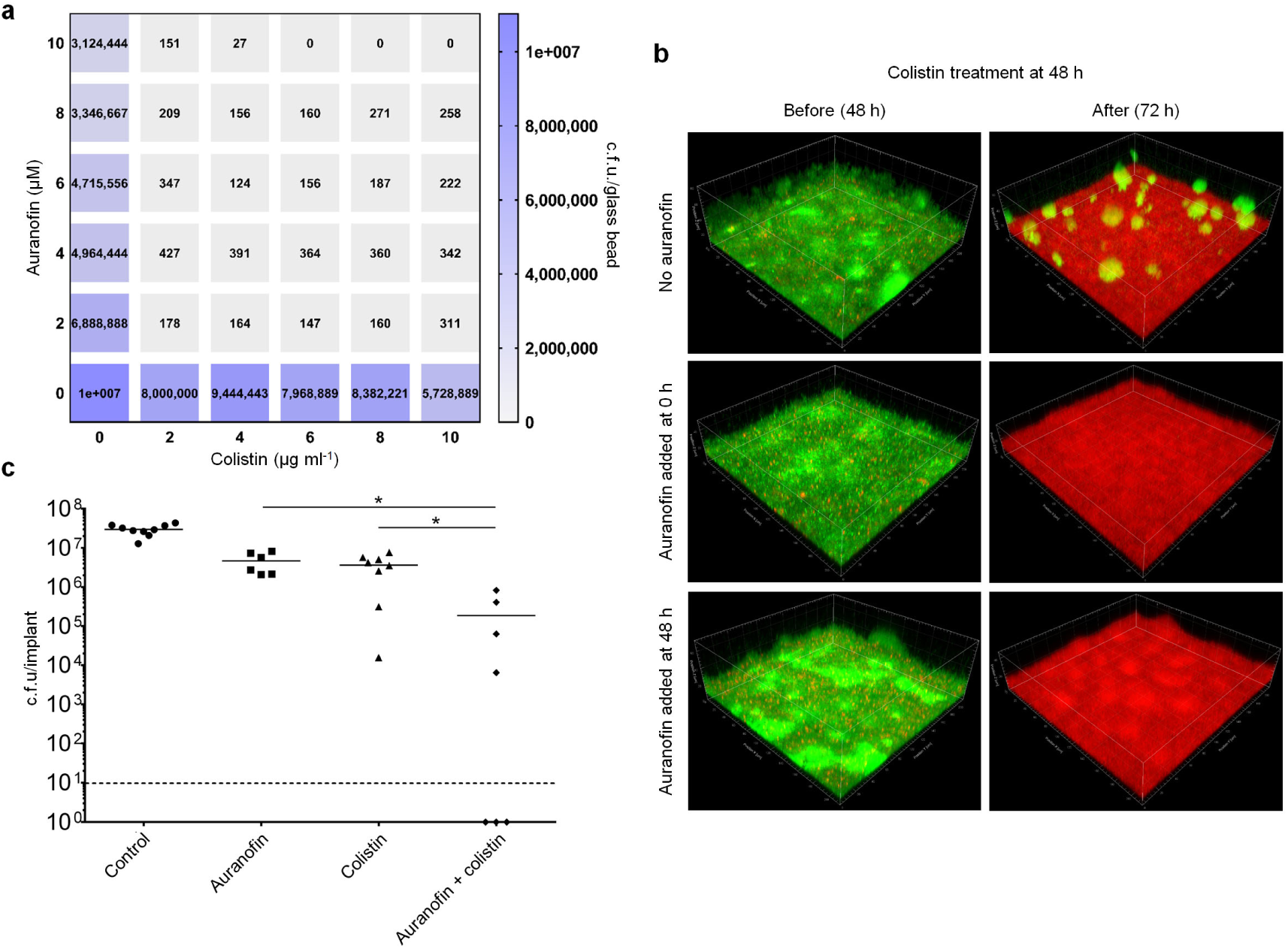
Auranofin displays strong synergy in eradicating *P. aeruginosa* biofilms when used in combination with colistin. (a) Mean c.f.u of cells isolated from one-day old PAO1 biofilms grown on glass beads before and after treatment with indicated combinations of auranofin and/or colistin in a checkerboard format. (b) Confocal fluorescence microscopy images of two-day old PAO1-*gfp* flow chamber biofilms before and after treatment with 10 µg ml^-1^ colistin. Auranofin (1 µM) was not added (top panels), added prior to biofilm formation (middle panels), or added together with colistin (bottom panels). Dead cells were revealed by staining with propidium iodide. (c) Mean c.f.u per implant of *in vivo* PAO1 biofilms obtained from silicone implants in the peritoneal space of mice, treated with auranofin (10 µM), colistin (1 mg kg^-1^) or a combination of both drugs. Dotted horizontal lines represent limit of detection. Mann-Whitney test: \**P* < 0.01.

To test whether auranofin and colistin can work together *in vivo*, we surgically inserted silicone implants coated with 1-day old *P. aeruginosa* biofilms into the peritoneal cavity of mice, and subsequently treated them locally with colistin (1 mg kg^-1^), auranofin (10 µM), or a combination of both drugs. Combination therapy of auranofin and colistin resulted in significantly lower bacterial titer counts from the implant than either auranofin or colistin alone (Fig. 6C). This confirms the synergy between auranofin and colistin in a murine model.

## Discussion

Auranofin is approved by the FDA for the treatment of rheumatoid arthritis. It has been shown to reduce inflammation of joints in rheumatic patients^44^; however, the precise mechanism of how this occurs is not clear. Auranofin also exhibits anti-cancer and anti-parasitic properties^39,45,46^. Furthermore, auranofin is able to kill a broad spectrum of Gram-positive bacteria, and also mycobacteria^40^. All of these activities may be attributed to auranofin’s ability to disrupt the thiol-redox balance in cells, in part by targeting the thioredoxin reductases in each of these systems^47^. Interestingly, auranofin is largely ineffective against most Gram-negative bacteria, except for a few species^48-50^, presumably because of lack of penetration across the bacterial outer membrane^51^. Here, we have demonstrated that auranofin is additionally able to inhibit virulence in *P. aeruginosa*. Despite not being able to accumulate at high concentrations in cells, auranofin causes downregulation of QS, TFP formation, and possibly T3SS. One possible target is Vfr, which controls all of these phenotypes. Consistently, we have shown that auranofin inhibits the ability of Vfr to bind to target promoter sequences in vitro. The fact that a cysteine-less Vfr variant is not inhibited in vitro indicates that auranofin also targets cysteines within this protein. Interestingly, *P. aeruginosa* does not gain resistance against auranofin even when expressing cysteine-less Vfr, suggesting that it has multiple targets in cells. This is probable, given that auranofin is a general thiol-targeting reagent^39^. Understanding the full mechanism of action of auranofin would require identifying these unknown targets.

Current clinical treatment options of *P. aeruginosa* infections involve the usage of carbapenems, aminoglycosides and ciprofloxacin, to name a few^16^. These drugs are often used in combination or singularly in treatment. Unfortunately, many strains of *P. aeruginosa* have developed resistance to such compounds and drastically reduce the pool of available treatment options for clinicians^7^. Colistin remains one of the few drugs effective against most strains of *P. aeruginosa*. Its usage, however, remains limited and is confined to last resort situations due to its toxicity^52^. We have shown that auranofin and colistin combine synergistically to completely eradicate *P. aeruginosa* biofilms, something not observed with colistin treatment alone. This synergy is also observed in a murine implant model. We believe that the strong synergy between auranofin and colistin works in both directions. It has been shown that disrupting outer membrane permeability (here with the use of colistin) allows auranofin to penetrate the cell envelope and kill Gram-negative bacteria^51^, through its potential action on thioredoxin reductase. At the same time, the inhibition of TFP and QS (by auranofin) during biofilm development may prevent the formation of colistin-tolerant subpopulations within the biofilm, thereby allowing complete eradication. The unique synergistic effects of auranofin and colistin may be exploited as a new treatment regimen for *P. aeruginosa* infections. Given that auranofin is already an FDA-approved drug, it may not be in the very far future that this approach can be tested.

## Materials and Methods

### Bacterial strains and growth conditions

All bacterial strains used in this study are summarized in Table S2. Strain PAO1 was used for all *in vivo* experiments^53^. High-throughput screening of quorum sensing inhibitors was done using strain PAO1-*lasB-gfp*(ASV)^34^. The extent of inhibition of QS was verified using strains PAO1-*P*_*rhlA*_-*gfp*(ASV) and PAO1-*P*_*pqsA*_-*gfp*(ASV)^35^. Luria-Bertani (LB) broth (1% tryptone and 0.5% yeast extract) and agar were prepared for growth of overnight cultures. M9 minimal media was prepared as previously described^54^. Glass bead biofilm and flow cell biofilm experiments were conducted in ABT minimal medium supplemented with 0.4 g l^-1^ glucose (ABTG)^43,55^. For plasmid maintenance in *Escherichia coli*, ampicillin (Amp) was used at 200 µg ml^-1^, tetracycline (Tc) at 12.5 µg ml^-1^. For marker selection in *P. aeruginosa*, gentamicin (Gm) was used at concentration 30 µg ml^-1^, tetracycline (Tc) at 50 µg m^-1^, carbenicillin (Carb) at 300 µg ml^-1^.

### Plasmid construction

All plasmids are listed in Table S3. To construct a plasmid with desired DNA fragment, gene was amplified by PCR using PAO1 genomic DNA template. Amplified DNA fragment was digested with appropriate restriction enzymes (New England Biolabs) and ligated into the same restriction sites of a plasmid. DH5α cells were transformed with the ligation product and selected on LB plates containing appropriate antibiotics. All constructs were verified by DNA sequencing (Axil Scientific, Singapore). To construct pME6031*vfr, vfr*, together with 200 bp upstream of the ATG translational start codon was amplified and inserted into the HindIII/SacI site of pME6031. To construct pUCP18vfr, *vfr* was amplified and inserted into the EcoRI/HindIII site of pUCP18. To construct pME6031*vfrΔ5* and pUCP18*vfrΔ5*, the 5 cysteine residues in *vfr* were sequentially replaced by serine residues via site-directed mutagenesis using C20S-FWD/C20S-REV, C38S-FWD/C38S-REV, C97S-FWD/C97S-REV, C156S-FWD/C156S-REV and C183S-FWD/C183S-REV primer pairs. For the above, pME6031*vfr* and pUCP18*vfr* were used as the templates, respectively. The resulting PCR products were digested with DpnI, and transformed into DH5α and selected on LB plates containing tetracycline and ampicillin, respectively.

To construct *P*_*lasR*_ and *rsaL* probes for EMSA, primer sequences are given in Table S4.

### QS Inhibitor Assay

The QS inhibitor assay was conducted as previously described^34^. Briefly, 5-ml overnight cultures were washed in M9 minimal medium and diluted to an OD_450_ of 0.2. 150 μl was added into a 96-well plate that contained 150 μl of each library compound in each well to appropriate concentrations. Compounds screened were from the triazine library^56^ (2,920 compounds) (Young-Tae Chang (POSTECH)) and a ENZO commercial library of FDA-approved drugs (640 compounds). The samples were incubated at 34 °C in a plate reader (Victor X2, Perkin Elmer). Green fluorescence at excitation wavelength of 490 nm was measured at 15 min intervals over 17 h and normalized to cell density at OD_450_.

### Quantitative Real-time Polymerase Chain Reaction (qRT-PCR)

The qRT-PCR experiment was conducted as previously described^57^. 5-ml cultures were grown (inoculated with overnight cultures at 1:100 dilution) in M9 minimal medium containing appropriate concentrations of auranofin/DMSO at 37 °C until OD_450_ reached ∼1.0. RNeasy Protect Bacteria Mini Kit with on-column DNase digestion (Qiagen) was used to extract total RNA. A Turbo DNA-free vigorous protocol was used for a second round of DNase treatment (Ambion). Agilent 2100 Bioanalyzer (Agilent Technologies) and Qubit 2.0 Fluorometer (Invitrogen) were used to assess the integrity of the total RNA and the level of DNA contamination. Ribosomal RNA contamination was removed using the Ribo-Zero Magnetic Kit (Bacteria) (Epicentre). Finally, RNeasy Mini Kit (Qiagen) with on-column DNase digestion was used to extract the total RNA. The purity and concentration of the RNA were determined by NanoDrop spectrophotometry, and the integrity of RNA was measured using an Agilent 2200 TapeStation System. The elimination of contaminating DNA was confirmed via the real-time PCR amplification of the *rpoD* gene using total RNA as the template.

Quantitative reverse transcriptase PCR (qRT–PCR) was performed using a two-step method. SuperScript III First-Strand Synthesis SuperMix kit (Cat. No. 18080-400, Invitrogen) was used to synthesize first-strand cDNA from total RNA as a template. The cDNA was used as a template for qRT–PCR using a SYBR Select Master Mix kit (Cat. No. 4472953, Applied Biosystems by Life Technologies) on an Applied Biosystems StepOnePlus Real-Time PCR System. *rpoD* was used as an endogenous housekeeping control. Melting curve analyses were employed to verify the specific single-product amplification.

### Elastase Assay

5-ml cultures were grown (inoculated with overnight cultures at 1:100 dilution) in M9 minimal medium containing appropriate concentrations of auranofin/DMSO at 37 °C until OD_450_ reached ∼1.0. Cells were then normalized according to OD_450_ and filtered. The elastase levels within the supernatant were measured using an Elastase Assay kit (EnzChek Elastase Assay kit, Life Technologies) according to manufacturer’s instructions.

### Measurement of Rhamnolipid Levels

5-ml cultures were grown (inoculated with overnight cultures at 1:100 dilution) in M9 minimal medium containing appropriate concentrations of auranofin/DMSO at 37 °C until OD_450_ reached ∼1.0. Cells were normalized according to OD_450_ and centrifuged at 4700 x *g* for 10 min. 380 μl of acetonitrile, 100 μl of 40 mM aqueous ammonium acetate and 20 μl of a 23 ppm acetonitrile solution of 16-hydroxydecanoic acid as an internal standard (IS) were added to 500 μl of filtered supernatant. 10 μl was subjected to LC-MS analysis.

The analysis of rhamnolipid (Rha-C10-C10) (Sigma-Aldrich) in culture samples was performed on an Ultra High Performance Liquid Chromatography system (UFPLC Ultimates 3000, Dionex, CA, USA) equipped with a C18 HPLC column (100 mm x 4.5 mm, 3.5 mm, 40 oC, Zorbax Plus) coupled with a quadrupole-ion trap mass spectrometer (QTRAPs 5500, AB Sciex, DC, USA). An acetonitrile-water gradient containing 4 mM of ammonium acetate was used. Chromatography was performed with initial 40% acetonitrile for 4 min and the acetonitrile concentration was raised to 90% after 20 min. The analytes were ionized in negative mode using electrospray ionization and analyzed using Multiple Reaction Mornitoring (MRM) on 4 mass transition pairs (Rha-C10-C10: 649.5>479.5, 649.5>169.5, 649.5>163; IS: 271.2>225.4). The concentration of Rha-C10-C10 in culture samples was quantified using a calibration curve plotted from six Rha-C10-C10 standard solutions ranging from (0 - 10ppm) spiked with a fixed concentration of 16-hydroxydecanoic acid as internal standard. All the standard solutions and samples were spiked with 16-hydroxydecanoic acid as an internal standard to improve accuracy of results and to remove experimental uncertainty from systematic and random errors. The chromatograms were integrated using Analyst 1.5.1 (AB Sciex, DC, USA).

### Measurement of pyocyanin levels

5-ml cultures were grown (inoculated with overnight cultures at 1:100 dilution) in M9 minimal medium containing appropriate concentrations of auranofin/DMSO at 37 °C until OD_450_ reached ∼1.0. Cells were then normalized according to OD_450_ and filtered. The absorbance of the supernatant was measured at 691 nm.

### Proteomics

5-ml cultures were grown (inoculated with overnight cultures at 1:100 dilution) in LB broth at 37 °C until OD_450_ reached ∼1.0. Cells were washed, re-suspended in fresh LB medium. After harvesting, the cell pellet was washed twice with 1X PBS and resuspended in 2 ml of lysis buffer containing 0.5 M triethylammonium bicarbonate (TEAB), 0.1 M SDS and protease inhibitor cocktails (Sigma-Aldrich). The cells were ruptured by probe sonication (5 min of sonication, 30% amplitude) in ice slurry and the cell debris was removed by centrifugation at 4 °C at 16000 x *g* for 10 min. The protein supernatant from three biological replicates for each growth condition were pooled together. 200 μg of proteins from each growth condition were dissolved in equal volume of Laemelli 2X sample buffer (Invitrogen, USA) supplemented with 0.5% 2-mercaptoethanol. The proteins were then denatured by boiling at 95 °C for 5 min. 2D-SDS-PAGE was performed using 10% separating gel to concentrate the protein samples, with a final 20% blocking gel which reduced protein loss by preventing the proteins from leaving the gel.

Gel fragments were submitted for iTRAQ proteomics analysis. Briefly, fragments were reduced with 5 mM tris 2-carboxyethyl phosphine hydrochloride (TCEP) followed by alkylation with 10 mM methyl methanethiosulfonate (MMTS). They were subsequently dehydrated with 100% acetonitrile (ACN) and digested using sequencing grade modified trypsin (Promega, USA) at 37 °C for 20 h. Peptides were dried using 50% ACN/5% acetic acid mixture and dried under vacuum.

iTRAQ labeling of peptide samples was performed using the iTRAQ reagent multiplex kit (Applied Biosystems, Life Technologies) according to manufacturer’s instructions. Both labelled samples were pooled and reconstituted in Buffer A (10 mM KH2PO4, 25% acetonitrile, pH 2.85), and fractionated using PolySULFOETHYL A SCX column by HPLC (Shimadzu, Japan) at a flow rate of 1.0 ml min^-1^. The collected fractions were desalted with Sep-Pak Vac C-18 cartridges (Waters, USA), dried under vacuum and reconstituted in 0.1% formic acid for LC-MS/MS analysis. LC-MS/MS analysis was conducted using a Q-Star Elite mass spectrometer (Applied Biosystems, USA) coupled to a micro flow HPLC system (Shimadzu, Japan).

### Immunoblot analysis

5-ml cultures were grown (inoculated with overnight cultures at 1:100 dilution) in LB broth at 37 °C until OD_450_ reached ∼1.0. Cells were normalized according to OD_450_ and mixed with equal amounts of 2X Laemmli reducing buffer and subjected to SDS-page on 2 separate gels. Protein bands for both of the gels were transferred to polyvinylidene fluoride (PVDF) membranes (Immun-Blot® 0.2 μm, Bio-Rad) using semi-dry electroblotting system (Trans-Blot® TurboTM Transfer System, Bio-Rad). Membranes were blocked by 1X casein blocking buffer (Sigma). α-Vfr was provided by Katrina Forest (University of Wisconsin-Madison) and used at a dilution of 1:25,000. α-rabbit IgG secondary antibody conjugated to HRP (from donkey) was purchased from GE Healthcare and used at 1:5,000 dilution. Luminata Forte Western HRP Substrate (Merck Milipore) was used to develop the membranes and protein bands were visualized by G:BOX Chemi XT 4 (Genesys version 1.3.4.0, Syngene).

### Motility assay

10-ml overnight cultures were grown in LB broth at 37 °C and normalized to the lowest cell density according to OD_450_. To determine swimming motility, 0.3% LB agar containing appropriate concentrations of auranofin/DMSO was point inoculated with the overnight into the middle and incubated at 30 °C for 24 h. To determine swarming motility, 0.5% LB agar containing appropriate concentrations of auranofin/DMSO was point inoculated with the overnight culture at the surface and incubated at 37 °C for 24 h. To determine twitching motility, 1% LB agar containing appropriate concentrations of auranofin/DMSO was point inoculated at the agar-plate interface with an overnight colony of plated cells and incubated at 37 °C for 48 h. The agar was subsequently removed and adhered cells were stained with 0.05% Coomassie blue G250 (Sigma) (40% methanol, 10% acetic). Plate images were visualized by G-BOX Chemi XT 4 (Genesys version 1.3.4.0, Syngene).

### Purification of Vfr and its variant

Purification of Vfr and VfrΔ5 was performed according to a procedure described previously with some modifications^58^. Vfr and VfrΔ5 were purified from pUCP-*vfr* and pUCP-*vfrΔ5*, respectively. For each strain, a 500-ml culture (inoculated from a mid-log culture at 1:100 dilution) was grown overnight in LB broth at 37 °C. Cells for each strain were harvested by centrifugation at 4,700 x *g* for 20 min. Cells were resuspended in 20 ml of 20 mM Tris.HCl, pH 8.0 containing 1 mM phenylmethylsulfonyl fluoride (PMSF), 100 μg ml^-1^ lysozyme and 50 μg ml^-1^ DNase I. The resuspended cells were lysed by a single passage through a high pressure French Press (French Press G-M, Glen Mills) homogenizer at 20,000 psi. Unbroken cells were removed by centrifugation at 5,000 x *g* for 10 min. The cell lysate was collected and centrifuged at 100,000 x *g* for 1 h in an ultracentrifuge (Model XL-90, Beckman Coulter). The supernatant was collected, loaded into a column packed with 1 ml of cyclic adenosine monophosphate (cAMP) agarose column (Sigma Aldrich), which had been pre-equilibrated with 10 ml of buffer A (25 mM Tris.HCl/150 mM NaCl, pH 8.0) and incubated at 4 °C for 1 h, by rocking. The resin mixture was later allowed to drain by gravity. The filtrate was collected, reloaded into column and drained as above. The column was washed with 4 x 10 ml buffer A and then eluted with 4 x 4 ml of buffer B (25 mM Tris.HCl/150 mM NaCl, pH 8.0) each with increasing amounts of free cAMP to 0.5, 5, 10 and 25 mM. The eluted fractions were concentrated in a 10 kDa cut-off ultra-filtration device (Amicon Ultra, Merck Millipore) by centrifugation at 4,000 x *g* to ∼200 μl. The concentrated sample was mixed with equal amounts of 2X Laemmli reducing buffer and subjected to SDS-PAGE analyses and immunoblotting. Fractions containing pure Vfr and VfrΔ5 were collected and used.

### Electromobility Shift Assay (EMSA)

0.5 μl of 4 mM auranofin was serial diluted in 10 μl of 1X Buffer C (20 mM Tris.HCl/200 mM KCl/2mM EDTA/1% glycerol/0.1 mg BSA, pH 8.0) and incubated with 4.8 μl of purified protein at appropriate concentration at room temperature for 10 min. The mixture was then combined with buffer D (1X Buffer C/500 μM cAMP/432 nM DNA probe/432 nM rsaL) and incubated for 20 min. 5 μl of 5X DNA loading dye (COMPOSITION) was added and the reaction was subjected to PAGE analyses on a 6% TBE gel (Tris.HCl/Boric acid/EDTA, pH8.0). DNA bands were stained by immersing the gel in 3X GelRed staining solution (3,000X GelRed stain/100mM NaCl) (Biotium) for 30 min followed by visualization by G-BOX Chemi XT 4 (Genesys version 1.3.4.0, Syngene)

### Glass beads biofilm assay

The glass bead biofilm assays were performed as described by Konrat, K. *et al*^59^. *Pseudomonas aeruginosa* PAO1 was cultured overnight in 2 ml LB broth at 37°C shaking condition (200 rpm). The overnight culture was centrifuged at 13,000 x *g* for 2 min and the supernatant was discarded. The cell pellet was washed once with 2 ml of 0.9% NaCl, followed by centrifugation to pellet the cells. After the wash, cells were then resuspended and adjusted to OD_600_ of 0.1 with ABTG before 1 ml of the culture was pipetted into each well of the 24-well, flat-bottom plate (Nunclon^®^ Δ Multidishes, ThermoFisher Scientific). One sterile 5-mm glass bead (Merck KGaA, Darmstadt, Germany) was placed into each well prior to incubation in 37 °C shaking condition (100 rpm) for 24 h for biofilm formation.

After 24 h of cultivation, the biofilm-coated glass beads were gently picked and submerged in 0.9% NaCl solution for 3 times using sterile tweezer to remove the unbound cells before transferring to new 24-well plates that consist of different concentrations of colistin sulfate (Sigma-Aldrich) and auranofin (Sigma-Aldrich) combinations. After 24 h of treatment in 37 °C shaking condition (100 rpm), each glass bead was then transferred to an 1.5-ml Eppendorf tube containing 1 ml of 0.9% NaCl before subjected to water-bath sonication using Elmasonic P120H (Elma, Germany; power=80% and frequency=37 KHz) for 10 min and rigorous vortex for 5 min to disrupt the biofilms. For enumeration of viable cells per glass bead, the samples were serially diluted, plated on LB agar followed by incubation overnight at 37 °C for colony forming unit (c.f.u.) count. Biological triplicates were performed and the average c.f.u. count per glass bead in each condition was determined.

### Dynamic development of *P. aeruginosa* biofilm in flow cell

The assembly of flow chamber and experimental setup has been described previously^43,57,60^. Bacterial strains were cultured overnight in 2 ml of LB medium in 37 °C shaking incubator (200 rpm). For marker selection in *P. aeruginosa*, 30 µg ml^-1^ gentamicin or 50 µg ml^-1^ tetracycline was used. The overnight cultures were adjusted to OD_600_ of 0.1 with ABTG and 500 µl of the diluted cultured was injected into each flow chamber and was incubated for 1 h without medium flow. Wild-type PAO1 were fed with ABTG containing either freshly prepared 2 µM auranofin or DMSO (as solvent control) at the rate of 4 ml h^-1^ via peristaltic pump (Cole-Parmer^®^) in 37 °C condition. QS-null strain PAO1 Δ*lasI*Δ*rhlI* was used as control. At time-point 6 h, 24 h, 48 h and 72 h, 1 flow cell from each condition was removed from the system and the bacterial cells were stained with 300 µl of 3.34 µM SYTO9 and 20 µM propidium iodide (PI) (LIVE/DEAD™ *Bac*Light™ Bacterial Viability Kit, Molecular Probes) for 15 min for the observation of live and dead cells respectively using confocal laser scanning microscopy. The experiments were performed in triplicate and representative images are shown.

### Auranofin and colistin treatment on *P. aeruginosa* flow cell biofilm

The flow cell biofilm experiments were setup as mentioned above. The GFP-tagged *P. aeruginosa* was cultured for 48 h prior to treatment with; DMSO (solvent control), or 1 µM auranofin single treatment, or 10 µg ml^-1^ colistin single treatment, or 1 µM auranofin and 10 µg ml^-1^ colistin combination treatment for 24 h. Then, the dead cells were stained with PI as described above prior to confocal microscopy imaging. The experiments were performed in triplicate and representative images are shown.

### Confocal microscopy imaging

The microscopy images of the biofilms were captured and acquired using LSM 780 confocal laser scanning microscopy (Carl Zeiss, Germany). Either 20x/0.80 DICII or 40x/1.30 DICIII (oil) objective lens was used for confocal imaging. 488 nm argon multiline laser and HeNe 561 nm laser were used to monitor the GFP-expressing or SYTO9-stained live cells and PI-stained dead cells respectively. The acquired images were further processed using IMARIS version 9 (Bitplane AG, Zurich, Switzerland).

### Image analysis (for biofilm development in flow cells)

The confocal images were processed using COMSTAT2 program to quantify the three-dimensional biofilm image stack as previously described^61-63^. The biofilm structures were described with 3 variables (biofilm biomass, mean thickness and roughness coefficient). All COMSTAT2 parameters are fixed at default setting prior to image analysis.

### Mouse implant infection model

All animal experiments were carried out in accordance to the NACLAR Guidelines and Animal and Birds (Care and Use of Animals for Scientific Purposes) Rules by Agri-Food & Authority of Singapore (AVA), with authorization and approval by the Institutional Care and Use Committee (IACUC) and Nanyang Technological University, under the permit number A-0191 AZ.

Seven to eight week-old female BALB/c mice (Taconic M&B A/S) were used in this experiment. The mouse implant infection model experimental setup was carried out as described previously^64,65^. *Pseudomonas aeruginosa* was cultured overnight in 2 ml of LB medium in 37 °C shaking incubator (200 rpm). The overnight culture was centrifuged at 13,000 x *g* for 1 min and the supernatant was discarded prior to washing with 0.9% NaCl at equal volume. The cell pellet was resuspended and diluted to OD_600_ of 0.1 with 0.9% NaCl. The biofilms were developed on cylindrical implants (4 mm × 5 mm Ø) in the diluted culture for 20 h at 37 °C with shaking condition (110 rpm). The implants were then washed 3 times with 0.9% NaCl to remove the unbound cells, before insertion into the peritoneal cavity of the mouse that was anesthetized with 100 mg kg^-1^ ketamine and 10 mg kg^-1^ xylene. 24 h after the insertion, 200 µl of the treatment; control (DMSO), or 10 µM auranofin, or 1 mg kg^-1^ colistin; or 10 µM auranofin and 1 mg kg^-1^ colistin was injected into the peritoneal cavity and the second treatment was given at 30 hours-post-infection. The mice were sacrificed at 48 hours-post-infection and the implants were retrieved, sonicated in a water-bath Elmasonic P120H (Elma, Germany; power=80% and frequency=37 KHz) for 10 min and vortexed rigorously 3 times of 10 seconds each, before subjected to c.f.u plating on LB agar plates to determine the viable bacterial cells on the implants. Experiments were performed with 10 replicates, and the results are shown as the mean. Implants from the mice that died before the experiment endpoint were not included in the statistical analysis.

## Acknowledgements

We thank Young-Tae Chang (POSTECH) for allowing access to small molecule libraries used in this study. We acknowledge Song Lin Chua (SCELSE) for his help in processing and submitting our samples for proteomics analysis. We thank Yingying Li (SCELSE) for her guidance in RT-PCR experiments. We are grateful to Katrina Forest (University of Wisconsin-Madison) for the generous gift of the α−Vfr antibody. This research was supported by the National Research Foundation and the Ministry of Education of Singapore under its Research Centre of Excellence Programme. M.G., S.-S.C., L.Y., L.Z.W.T., and Z.H. have filed an international patent application (PCT/SG2016/050436, filed 6^th^ September 2016) and granted a US patent (US 10,258,640 B2, dated 16^th^ April 2019) on the work described in this manuscript.

## Author Contributions

L.Y., M.G., and S.-S.C. conceived the project and directed the study. Z.H. conducted the QS inhibitor screening assay. L.Z.W.T. conducted all biochemical experiments. J.K.H.Y. and M.M.S.S. performed the mouse model and the biofilm experiments. B.Y.W. conducted the rhamnolipid analysis with supervision from S.F.Y.L.. L.Z.W.T., J.K.H.Y, and S.-S.C. wrote the manuscript with input from all authors.

