## Supporting information for "Auranofin inhibits virulence in *Pseudomonas aeruginosa*"

**Table S1. Fold change of proteins regulated in the presence of auranofin.**

| | Protein Name | Fold change (p<0.05) | | Protein Name | Fold change (p<0.05) |
|---|---|---|---|---|---|
| Efflux pump[a] | MexP | 47.8 | QS | PqsE | -5.0 |
| | MexQ | 44.0 | | PqsD | -4.2 |
| | OmpE | 45.2 | | PqsA | -4.2 |
| | MexE | 25.8 | | PqsC | -3.8 |
| | MexF | 25.3 | | PqsL | -3.7 |
| | MexB | 1.8 | | PhnA | -3.3 |
| | OprM | 2.3 | | RhlA | -3.0 |
| Porin[b] | OprC | -4.6 | | PqsH | -2.7 |
| | OprE | -2.9 | | PvdL | -1.5 |
| | OprD | -1.6 | | RhlR | -1.5 |
| QS[c] | PhzF1 | -47.9 | | PvdF | -1.4 |
| | PhzE1 | -19.6 | T3SS[d] | ExoS | -6.6 |
| | ChiC | -13.2 | | PopD | -5.0 |
| | PhzS | -13.1 | | PscQ | -4.0 |
| | PhzG1 | -11.3 | | ExsD | -3.1 |
| | PhzD1 | -10.7 | | ExsA | -2.4 |
| | RhlB | -6.1 | TFP[e] | PilM | -1.5 |
| | CbpD | -5.7 | | PilY | -1.4 |

[a] In the efflux pump category, MexA, MexT and OprN are also upregulated. However, due to the low number of peptides detected for these proteins, the changes are not considered significant.

[b] In the porin category, OprB and OprF are also downregulated. However, due to the low number of peptides detected for these proteins, the changes are not considered significant.

[c] In the quorum sensing category, LasR, PqsB, PvdJ, PvdO, PvdR, PhzB1, PhzC1, PhnB and PhzM are also downregulated. However, due to the low number of peptides detected for these proteins, the changes are not considered significant.

[d] In the T3SS category, ExoT, ExsC, PscC, PscD, PscJ, PscK, PopB, PopN and PcrV are also downregulated. However, due to the low number of peptides detected for these proteins, the changes are not considered significant.

[e] In the TFP category, PilA, PilC, PilG, PilN, PilO, PilW and PilX, are also downregulated. However, due to the low number of peptides detected for these proteins, the changes are not considered significant.

**Table S2. Bacteria strains used in this study.**

| Strains | Relevant genotypes or characteristics | Reference |
|---|---|---|
| PAO1 | Wild-type *P. aeruginosa* | 1 |
| NovaBlue | *endA1 hsdR17* ($r_{K12}^{-}$ $m_{K12}^{+}$) *supE44 thi-1 recA1 gyrA96 relA1 lac* F′ [*proA*$^{+}$*B*$^{+}$*lacI*$^{q}$*Z*Δ*M15*::Tn*10*] | Novagen |
| BL21(λDE3) | *F*$^{-}$ *dcm ompT hsdS*($r^{-}_{B}m^{-}_{B}$) *gal* (*λ*DE3) | Novagen |
| DH5α | F$^{-}$, ø80d*lacZ*ΔM15, Δ(*lacZYA-argF*) U169, *deo*R, *recA*1, *endA*1, *hsdR*17(rK$^{-}$, mK$^{+}$), *phoA*, *supE*44, λ–, *thi*-1, *gyrA*96, *relA*1 | 2 |
| PAO1-*lasB*-*gfp*(ASV) | PAO1 containing the *lasB*-*gfp* translational fusion; Gm$^{R}$ | 3 |
| PAO1-*rhlA*-*gfp*(ASV) | PAO1 containing the *rhlA*-*gfp* transcriptional fusion; Gm$^{R}$ | 4 |
| PAO1-*pqsA*-*gfp*(ASV) | PAO1 containing the *pqsA*-*gfp* transcriptional fusion; Gm$^{R}$ | 4 |
| PAO1 Δ*vfr* | Virulence defective *vfr* mutant in PAO1 | This study |
| PAO1 Δ*lasI*Δ*rhlI* | *las* and *rhl* double mutant in PAO1; Gm$^{R}$; Tc$^{R}$ | 5 |
| PAO1-*gfp* | PAO1 expressing the green fluorescent protein | 6 |

**Table S3. Plasmids used in this study.**

| Plasmids | Relevant genotypes or characteristics | References |
|---|---|---|
| pME6031 | Lower copy, broad host-range cloning vector which is maintained in Gram-negative bacteria without selection pressure, $Tc^R$ | 7 |
| pUCP18 | High copy *P. aeruginosa*/*E. coli* shuttle vector $Amp^R$/$Carb^R$ | 8 |
| pME6031*vfr* | Encodes full length Vfr under its native promoter; $Tc^R$(p-*vfr*) | This study |
| pME6031*vfr*Δ*5* | Encodes full length VfrΔ5 under its native promoter; $Tc^R$(p-*vfr*Δ5) | This study |
| pUCP18*vfr* | Encodes full length Vfr under its native promoter; $Amp^R$/$Carb^R$ (pUCP-*vfr*) | This study |
| pUCP18*vfr*Δ*5* | Encodes full length VfrΔ5 under its native promoter; $Amp^R$/$Carb^R$ (pUCP-*vfr*Δ5) | This study |

**Table S4. Primers used in this study.**

| **Primers** | **Sequence (5' to 3')*** |
|---|---|
| Vfr-N-NdeI-FWD | ATGC CATATG GTA GCT ATT ACC CAC ACA CCC |
| Vfr-C-XhoI-REV | ATGC CTCGAG GCG GGT GCC GAA GAC |
| Vfr-N-HindIII-FWD | ATCG AAGCTT GAT AGC TGC GTC GCA AAA |
| Vfr-C-SacI-REV | ATCG GAGCTC TCA GCG GGT GCC GAA GAC |
| Vfr-N-EcoRI-FWD | ATCG GAATTC ATG GTA GCT ATT ACC CAC ACA C |
| Vfr-C-HindIII-REV | ATCG AAGCTT TCA GCG GGT GCC GAA GAC |
| $P_{lasR}$-FWD | TGC GCC TCG GAT CGC CCG GCC GAG |
| $P_{lasR}$-REV | TGT GCC GGA AAG GCA CAT TTC AGT C |
| *rsaL*-FWD | ATG CGA GTG CGT CAT AAC CAT CGA T |
| *rsaL*-REV | ATG CTT GGG GCT GTG TTC TCT C |
| Vfr C20S-FWD | GCA CAC AGC CAC CGC CGC |
| Vfr C20S-REV | GCG GCG GTG GCT GTG TGC |
| Vfr C38S-FWD | GGC GAT CGC AGC GAA ACG CTG TTC |
| Vfr C38S-REV | GAA CAG CGT TTC GCT GCG ATC GCC |
| Vfr C97S-FWD | GCC AAG GTG GAA AGC GAA GTC GCC |
| Vfr C97S-REV | GGC GAC TTC GCT TTC CAC CTT GGC |
| Vfr C156S-FWD | CTG GAC CTG AGC CAG CAA CCG GAC |
| Vfr C156S-REV | GTC CGG TTG CTG GCT CAG GTC CAG |
| Vfr C183S-FWD | ATC GTC GGC AGC TCG CGG GAA |
| Vfr C183S-REV | TTC CCG CGA GCT GCC GAC GAT |

*Relevant restriction sites are underlined.

### Figure S1

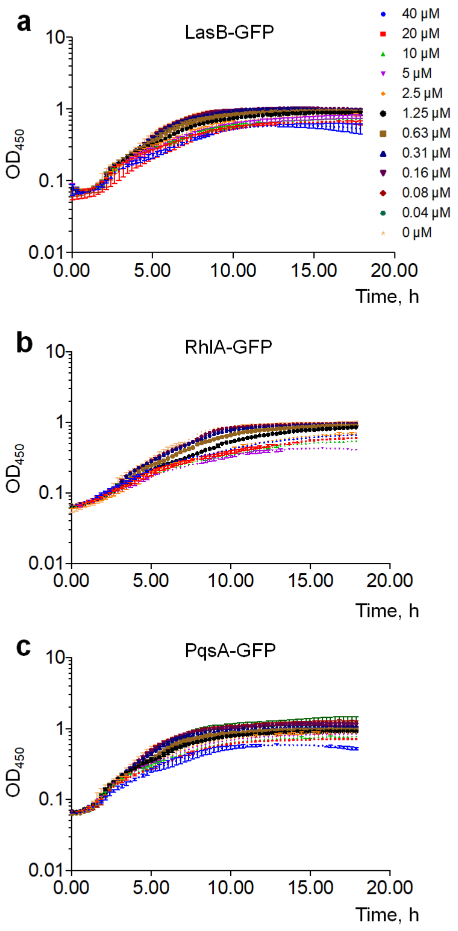


**Figure S1.** Growth profiles of screening strains are minimally affected by auranofin. Cell density, measured at 450 nm, for (a) PAO1-*lasB*-*gfp*, (b) PAO1-$P_{rhlA}$-*gfp*, and (c) PAO1-$P_{pqsA}$-*gfp* strains, expressed as a logarithmic curve against time.

**Figure S2**

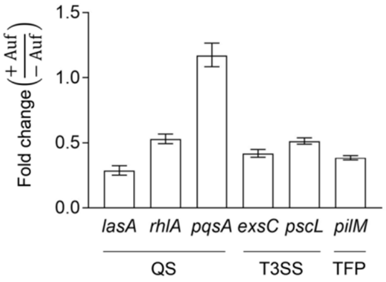


**Figure S2.** The cellular transcript levels of key QS, T3SS and TFP genes are reduced in the presence of auranofin. The RNA levels of indicated genes in the presence of auranofin are expressed as fold change compared to that in the absence of auranofin. Error bars represent standard deviation from biological triplicates. The lack of any apparent effect of auranofin on *pqsA* transcript levels was unexpected but may be related to sampling timing not well-aligned with the period of maximal expression (late stationary phase).

**Figure S3**

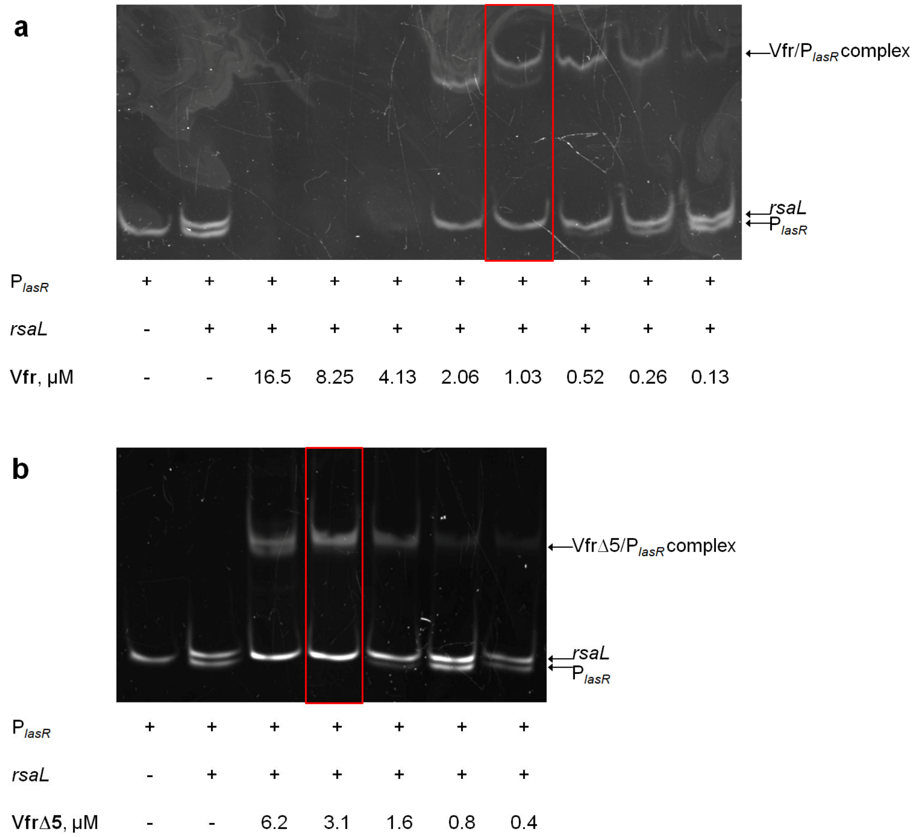


**Figure S3.** Purified Vfr binds specifically to target promoter sequences *in vitro*. Representative polyacrylamide gels showing (a) Vfr and (b) VfrΔ5 binding to $P_{lasR}$ at indicated Vfr protein concentrations. In these assays, Vfr and VfrΔ5 bind specifically to target the promoter sequence at < 1 and 3 µM, respectively. The coding region of the gene *rsaL* is used as a probe to control for non-specific binding.

### Figure S4

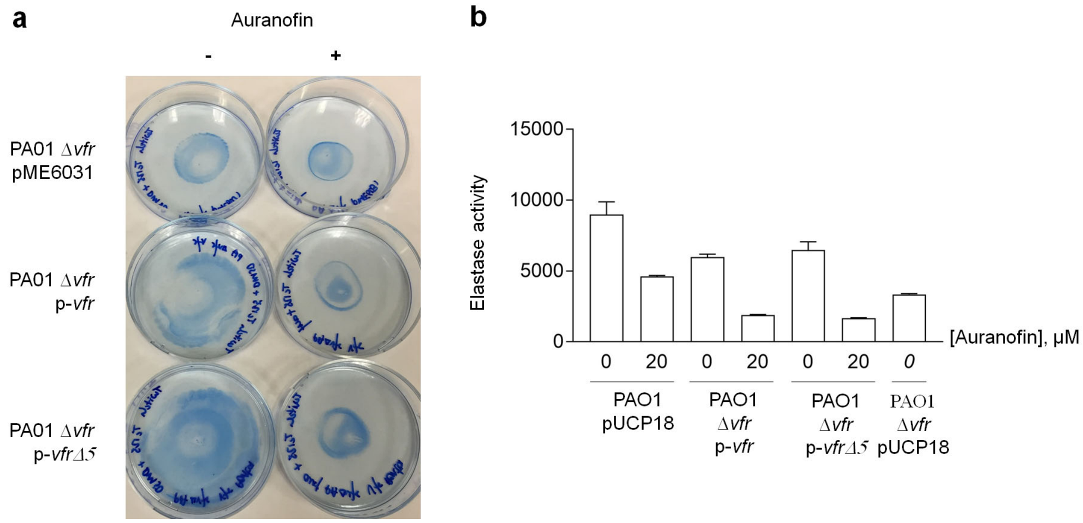


**Figure S4.** VfrΔ5 is able to restore Vfr-controlled phenotypes in Δ*vfr* cells, but these phenotypes can still be inhibited by auranofin. (a) TFP-dependent twitching (1% LB agar) motility of Δ*vfr*, or complemented strains in the presence or absence of 20 μM auranofin. Cells at the base of the plates were stained with Coomassie blue. (b) QS-controlled elastase activity in the supernatants from cultures of Δ*vfr*, or complemented strains in the presence or absence of 20 μM auranofin. Error bars represent standard deviation from biological triplicates.

**Figure S5**

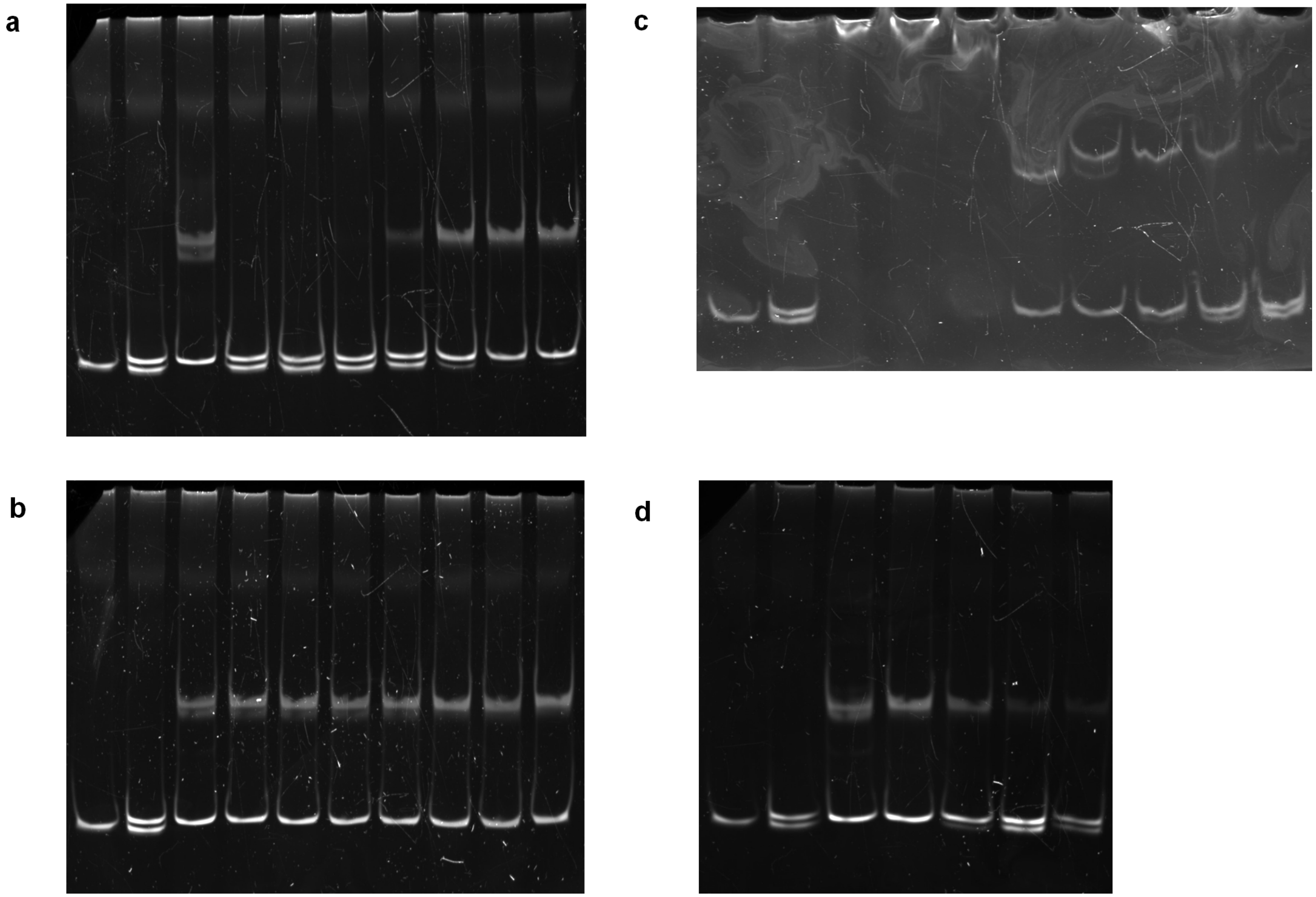


**Figure S5.** Uncropped gels for (a) Fig. 4A, (b) Fig. 4B, (c) Fig. S3A, and (d) Fig. S3B.
